# The BHMT-TET1 axis regulates glycolytic metabolism in oligodendrocytes and increases myelin in the EAE mouse model of multiple sclerosis

**DOI:** 10.64898/2026.07.22.737014

**Authors:** Marwan Shalih Maraicar, Sarah Sternbach, Morgan W. Psenicka, Katherine Knies, Ethan Lesco, Najmah Al Ramel, Andrew Eagar, Steven Zeisel, Ernest J. Freeman, Robert Clements, Jessica L. Williams, Jennifer McDonough

## Abstract

The inability of oligodendrocyte progenitor cells (OPCs) to mature into myelin-making oligodendrocytes (OLs) is a major contributor to disease and disability in multiple sclerosis (MS). Oligodendrocyte maturation is a tightly controlled process with a strong reliance on epigenetic regulation involving DNA methylation and hydroxymethylation. We have previously shown that one carbon metabolism is dysregulated in MS, specifically the methyl donor betaine is depleted in the MS brain. Betaine donates methyl groups to betaine homocysteine methyltransferase (BHMT) in the methionine cycle to increase S-adenosylmethionine (SAM) for epigenetic methylation processes. In the present study we tested the effects of activating the BHMT methylation pathway on preventing MS pathology. We describe a novel mechanism mediated by BHMT and the Ten-eleven translocator enzyme (TET1) that converts 5-methylcytosine (5-mC) to 5-hydroxymethylcytosine (5-hmC). We show that this pathway supports oligodendrocyte metabolism to enhance myelin and reduce clinical disability in the experimental autoimmune encephalomyelitis (EAE) mouse model of MS. ChIP-seq studies show that BHMT is enriched at genes involved in OPC metabolism and proximal ligation assays (PLAs) demonstrate that BHMT interacts with TET1 on chromatin. This interaction regulates gene expression programs that support a shift in OPC metabolism to glycolysis during neuroinflammatory processes. These data highlight the critical role of methionine metabolism in supporting myelination and have important implications for the development of new therapeutic strategies for MS and other neurodegenerative diseases.

## 1. Introduction

Multiple sclerosis (MS) is an inflammatory and neurodegenerative disease of the central nervous system accompanied by oligodendrocyte cell death and demyelination. Disease modifying therapies for MS include ocrelizumab and copaxone that act to suppress peripheral B and T cells and limit inflammatory demyelination. While these therapies have been effective in preventing relapses in MS, there still is no cure. The experimental autoimmune encephalomyelitis (EAE) mouse model of MS has been helpful in identifying additional therapies that suppress damaging peripheral immune T-cell activation (Al-Mazroua et al., 2022; Alghibiwi et al., 2023), however therapies that can stimulate remyelination are still needed. The oxidative environment in MS renders OPCs unable to differentiate into myelin-making mature OLs increasing the incidence and severity of white matter cortical lesions and neurological disability (Kuhlmann et al, 2008; Starost et al, 2020). It has been well established that OPC maturation is epigenetically regulated by programs of transcriptional activation and repression mediated by histone and DNA methylation (Hernandez & Casaccia, 2015; Liu et al., 2015, 2016; Yen et al, 2016; Moyon et al., 2017). While DNA methylation silences gene expression, the Ten eleven translocator 1 (TET1) mediated conversion of 5-methylcytosine (5-mC) to 5-hydroxymethylcytosine (5-hmC) can activate gene expression and has been shown to enhance remyelination (Moyon et al., 2021).

It has been shown that a dysfunction of mitochondria is involved in neurodegenerative processes in MS (Dutta et al., 2006) and therapies that can restore mitochondria are being actively pursued (Mozafari et al., 2025). During differentiation of OPCs to myelinating OLs there is a shift in energy metabolism to increased glycolysis. As OPCs mature, they rely increasingly on glycolytic energy metabolism (Funfschilling et al., 2012). This is believed to be due to the increase in acetyl-CoA required to synthesize myelin lipids that can be generated from glycolysis. The gene expression changes that mediate this change in metabolism are largely unknown; however, it has been clearly established that differentiation of OPCs is controlled by transcriptional activation and repression processes that are influenced by epigenetic modifications. Much of this epigenetic regulation relies on one carbon metabolism for the synthesis of S-adenosylmethionine (SAM), the methyl donor required for histone and DNA methylation (Obeid et al., 2013). In MS and other oxidatively stressful conditions, the one carbon cycle enzymes and metabolites are downregulated, including SAM, and subsequent gene transcription of energy metabolism genes is abnormal (Singhal et al., 2015; 2018). We have previously shown that the methyl donor betaine restores the one carbon cycle under oxidative conditions enhancing methylation to restore epigenetic control of energy metabolism (Singhal et al, 2020; Sternbach et al, 2021). Enhancing one carbon metabolism with dietary factors such as methionine or betaine, can alter levels of the methyl donor SAM and influence methyltransferase activity and gene expression. This is due to the fact that the Km of many methyltransferases overlap with SAM concentrations in cells (Mentch et al., 2015). Other studies have shown that betaine has anti-aging effects and supplementation exerts prominent anti-inflammatory effects in the nervous system and can alter NMDA signaling (Zhang et al., 1996; Pummer et al., 2000; Ortiz-Costa et al., 2008; Yang et al., 2018; Chang et al., 2025). Betaine action to enhance methionine metabolism and levels of the methyl donor SAM, requires the enzyme BHMT. Prieur et al. (2017) showed that the absence of BHMT *in vivo* resulted in decreased total brain volume and impaired short term and reference memory.

This study seeks to clarify the impairment of energy metabolism and maturation in CNS myelinating cells and their precursors, focusing particularly through the lens of one carbon metabolite manipulation to epigenetically influence gene transcription. We have previously demonstrated in neurons that the BHMT enzyme binds to and activates gene expression vital for mitochondrial function and that betaine can ameliorate symptoms in the cuprizone mouse model of MS, but the specific role of this pathway in oligodendrocytes remains unclear (Singhal et al, 2015; Singhal et al, 2020). In the present study we show that OPC metabolism is shifted to glycolysis by betaine and this switch is dependent on BHMT and TET1. In the EAE mouse model of MS, betaine administration leads to an increase in myelin and prevents neurodegeneration, suggesting that betaine is neuroprotective and may support remyelination.

## 2. Materials and Methods

### 2.1 EAE mice

All procedures were approved by the Kent State University Institutional Animal Care and Use Committee (IACUC) (Animal Care Assurance number D16-00344). Twenty C57Bl/6J wild type (WT) mice were obtained commercially from The Jackson Laboratory (RRID: IMSR_JAX: 000664) and housed under specific pathogen-free conditions. Animals of mixed sex (10 males, 10 females) were arbitrarily selected into control or EAE groups at 10weeks of age. Mice were immunized subcutaneously with 100μl of a standard emulsion (Hooke Laboratories, EK2110) containing complete Freund’s adjuvant and MOG_35-55_ both below the scruff and on the lower back. Pertussis toxin (80ng) (Hooke Laboratories, EK-2110) was injected intraperitoneally (i.p.) on the day of immunization and 48 hours later. Mice were monitored daily for clinical signs of disease as follows: 0, no observable signs; 1, limp tail; 2, limp tail and ataxia; 2.5, limp tail and knuckling of at least one limb; 3, paralysis of one limb; 3.5, partial paralysis of one limb and complete paralysis of the other; 4, complete hindlimb paralysis; 4.5, moribund; 5, death. At day 15 post injection, corresponding to peak of clinical disability, 10 mice were given i.p. injections of betaine (50 mg/day) and 10 with vehicle (PBS) every day until day 40 when they were sacrificed. This level of betaine is based on previous studies that have shown that these levels of betaine influence epigenetic regulation (Chai et al., 2013; Singhal et al., 2020). The daily administration of betaine is also consistent with pharmacokinetics of betaine showing that the elimination half-life is over 14 hrs (Schwahn et al., 2003). For euthanizing mice, they were first anesthetized by inhalation of 3-5% Isoflurane (2-chloro-2-(difluoromethoxy)-1,1,1-trifluoro-ethane) in oxygen and then perfused with PBS followed by 4% paraformaldehyde (PFA) (Sigma, P6148). Significant differences in clinical disability between EAE groups were determined by a Mann-Whitney test, p < 0.05 considered significant.

### 2.2 Immunohistochemistry

For immunohistochemistry (IHC), mice were anesthetized by inhalation of 3-5% isoflurane and then intracardially perfused with PBS followed by 4% paraformaldehyde (PFA), and CNS tissue was removed and fixed in 4% PFA at 4°C for 24 h. Tissue was then cryopreserved in 30% sucrose (Millipore, 107651) frozen in O.C.T. Compound (Fisher HealthCare, 23-730-571). Frozen, sequential transverse sections (11 µM) were slide mounted and stored at − 80°C. Tissue sections were blocked with 10% goat serum and 0.1% Triton X-100 (Southern Biotech) for 1h at room temperature and then incubated with anti-myelin basic protein (MBP) (Abcam; ab7349; RRID: AB_305869) or NeuN primary antibody (EMD Millipore; ABN91) overnight at 4°C. Secondary antibody conjugated to Alexa Fluor 488 or Alexa Fluor 555 (ThermoFisher Scientific; RRID: AB_2534074) was applied for 1h at room temperature. Nuclei were counterstained with DAPI (ThermoFisher Scientific; RRID: AB_2307445) diluted in PBS. Sections were analyzed using the 20× objective of a confocal microscope LSM 800 (Carl Zeiss; RRID: SCR_023607). For IHC experiments, quantitation was performed from of 4–8 images taken across three to four lumbar spinal cord tissue sections at least 100μm apart per individual mouse. For MBP quantitation was done from four EAE mice treated with vehicle and six EAE mice treated with betaine. For NeuN, quantitation was done from three control mice, three controls treated with betaine, four EAE mice, and four EAE mice treated with betaine. The mean positive area, intensity, and Mander’s coefficient of colocalization were determined by setting thresholds using appropriate isotype control antibodies and quantified using ImageJ software (NIH) (RRID: SCR_003070). Statistical significance was determined in GraphPad Prism with a two tailed t-test, p ≤ 0.05 considered significant.

### 2.3 CARS microscopy to measure myelin lipids

Quantitative analysis of the myelin sheath lipid composition was measured in spinal cords from EAE mice treated with betaine or vehicle using Coherent Anti-Stokes Raman Scattering Microscopy (CARS) with the STELLARIS 8 CRS system (Leica Microsystems; RRID: SCR_008960). Initially, the spinal cords were fixed in a 4% paraformaldehyde solution for 24 hours. Subsequently, they were incubated in a 30% sucrose solution for a duration of one week. The lumbar region of the frozen spinal cord embedded in OCT compound was cryosectioned into 30μm sections, which were then fixed onto poly-L-lysine coated slides. These unstained sections were mounted in distilled water and cover-slipped, making them suitable for CARS imaging. The CARS imaging was conducted using a Leica STELLARIS 8 CRS system equipped with a 40x/1.10 water immersion objective and an oil immersion condenser with a numerical aperture (NA) of 1.1 (RRDI: SCR_024664). To capture CARS signals, a CARS 2000s Wheelfilter and a TransPMT Detector were employed. The Stokes laser was set at a wavelength of 1031.3 nm with a power output of 300 mV, while the pump laser operated at 797.9 nm with a power output of 150 mV. These settings correspond to the resonant frequency of 2855 cm^-^¹, which is characteristic of the CH vibrational signals of lipids. Both CARS imaging for lipids and Second Harmonic Generation (SHG) for collagen were acquired simultaneously, with a scanning speed of 200 Hz in a bidirectional manner. The pixel size was set to 0.284 μm, with a pixel dwell time of 4.175 microseconds. Images were acquired from lumbar spinal cords from six EAE mice treated with betaine or treated with vehicle (PBS) beginning at day 15 post injection and continuing until day 40 when mice were sacrificed. The acquired images were subjected to post-processing and analysis using FIJI-ImageJ software. The Corrected Total Cell Fluorescence (CTCF) was calculated to quantify the intensity of CARS signals associated with lipids in the myelin sheath. This was achieved by subtracting the Integrated Density values from the product of the Area of the Region of Interest (ROI) and the Mean Fluorescence of the Background. Statistical significance of lipid fluorescence in lumbar spinal cord between EAE mice treated with PBS and EAE mice treated with betaine was determined in GraphPad Prism with a two-tailed t-test, p < 0.05 was considered significant.

### 2.4 Proximity ligation assay

To determine whether BHMT interacts with TET1 on chromatin, a Proximity Ligation Assay (PLA) was performed using the Duolink PLA Kit from Millipore Sigma (DUO92101) according to manufacturer’s instructions. This assay detects protein interactions or protein-DNA interactions within 40 nm of each other. Initially, WT C57Bl/6 mouse spinal cords were fixed in a 4% Paraformaldehyde solution for 24 hours. Subsequently, they were incubated in a 30% sucrose solution for a duration of one week. The lumbar region of the frozen spinal cord was cryosectioned into 30 μm sections, which were then fixed onto poly-L-lysine coated slides. Primary antibodies raised in different species, BHMT and TET1 or BHMT and 5-hmC (RRID: AB_2274828; RRID: AB_3262620; RRID: AB_10013602), were used at 1:500 dilution. Once the antibody solutions were added to the slides, the samples were incubated at 4°C for 24 hours. They were then incubated in PLUS and MINUS PLA probes specific to the corresponding primary antibody species. Ligase was added with ligation buffer at 1:40 dilution followed by DNA Polymerase to amplify the PLA probes. For a negative control, the primary antibodies were excluded for some sections. These were then incubated with only the PLA secondaries. The slides were mounted with a coverslip using mounting medium with DAPI. Images were acquired using Confocal microscopy (Olympus FV100) (RRID: SCR_020337) with a 60x objective at λex 495 nm; λem 527 nm (green) for the PLA signals and λex 358 nm; λem 461 nm (Blue) for DAPI. The acquired images were subjected to post-processing and analysis using FIJI-ImageJ software. The figures were represented as Maximum Intensity Projections with a Z-Stack size of 30.

### 2.5 Oligodendrocyte isolation and culture

For isolating primary OPC cultures, C57BL/6 wild type (WT) mice and BHMT^-/-^ mice were timed-bred overnight in the vivarium (12L:12D cycle) with access to food and water *ad libitum*. WT mice were purchased from Jackson Laboratories (000664). BHMT^-/-^ mice exhibit a constitutive knockout of BHMT throughout the body (Teng et al., 2011). These mice were a gift from Dr. Steven Zeisel at UNC. In the morning, males were separated and females with a sperm plug were noted as embryonic day 0.5 (E 0.5). Primary OPCs were isolated from pregnant C57BL/6 WT and BHMT^-/-^ mice at E18 following an established protocol (Yang et al, 2016). In brief, pregnant dams were euthanized with carbon dioxide followed immediately by a midline abdominal incision opening of the uterus and extraction of the pups. The E18 pups were removed from the pregnant dams and decapitated. Brains were removed and placed in cold 1x PBS, and cortices and meninges were further dissected out. Cortices were moved to new culture dish and chopped with a blade, followed by trituration with growth medium to homogenize the tissue. The tissue was then collected and plated in a T-25 flask with growth medium with a media change after 72 hours. Media was then changed every other day for 10 days, and flasks were then placed on an orbital shaker overnight (37 °C, 250 rpm). The next day, media was removed from the flasks, strained through a 40 µm filter, and centrifuged at 900 rpm for 10 minutes. The pellet was resuspended in OPC basal medium and incubated in non-coated culture dish at 37 °C for 45 minutes to allow differential adhesion of astrocytes and microglia. Cells were then collected and replated in PDL-coated plates at the appropriate density and left to grow for 7 days. Approximately thirty timed pregnant rats (about 250 E18 embryos) were required to obtain OPCs for ChIP-seq, qRT-PCR, DNMT assays, and respirometry experiments as described.

### 2.6 ChIP-seq

To explore BHMT binding patterns, ChIP-seq was performed with chromatin isolated from primary mouse OPCs and an antibody to BHMT (ProteinTech; RRID: AB_2290472). Using a ChIP-IT Express Kit (ActiveMotif 53008), cells were fixed with 37% formaldehyde in base medium for 10 min. Fixation was stopped by the addition of a 1x glycine stop-fix solution in PBS for 5 min, followed by scraping with a rubber policeman to collect the fixed cells. The cells were then lysed in a Dounce homogenizer and centrifuged at 2400xg at 4°C for 10 min to pellet the nuclei. The pellet was then resuspended in shearing buffer with protease inhibitors (Sigma, P8340) and sheared by sonication with a QSonica sonicator for 25 pulses, 30 sec on and 30 sec off. Resulting fragments were 200-300 bp in length as assessed by electrophoresis. With 10 µl of sheared sample taken out as a control, the IP reaction was set up sequentially: 25 µl protein A/G magnetic beads, 10 µl of ChIP buffer, 22 µg of sheared chromatin, 1 µl of protease inhibitors, 3 µg of BHMT antibody and distilled water to make the final volume up to 100 µl. The IP reaction was incubated overnight on an end-over-end rotator at 4°C. Beads were separated with a magnetic stand and washed with ChIP buffer before eluting the chromatin. Chromatin was cleaned with proteinase K treatment and stored at −20 °C. Library preparation, sequencing, and analysis was performed at Novogene.

Enriched peaks were identified and isolated using MACS14 (Zhang et al. 2008) (RRID: SCR_013291) on the Docker container platform (Merkel 2014). Immunoprecipitated sequences were compared against untreated reference mouse OPCs and the *Mus musculus* genome using a p-value threshold of < 0.001. Overlaps between transcription start sites (TSS) and enriched peak regions within 5kb of a TSS were analyzed in R version 4.2.1 (RRID: SCR_001905) (R Core Team 2022) using Bioconductor version 3.14 (RRID: SCR_006442) (Gentleman et al. 2004, Huber et al. 2015) and the following R packages: GenomicRanges and IRanges (RRID: SCR_000025, RRID: SCR_006420) (Lawrence et al. 2013), rtracklayers (RRID: SCR 021325) (Lawrence et al. 2009), chipseq (Sarkar et al. 2021), Gviz (RRID: SCR_024239) (Hahne & Ivanek 2016), biomaRt (RRID: SCR_019214) (Durinck et al. 2005, 2009), and BSgenome.Mmusculus.UCSC.mm10 (The Bioconductor Dev Team 2014).

### 2.7 qRT-PCR

To confirm the functional implications of our ChIP-seq data on glycolysis, qRT-PCT was performed on OPCs treated overnight with 100 µM sodium nitroprusside (SNP) (Sigma-Aldrich, 71778) and/or 1 mM betaine (Thermo Scientific, B24397.36) with the Agilent SYBR Green qRT-PCR kit (Agilent, 600886). Cells were treated with SNP to mimic oxidative and inflammatory conditions observed in MS resulting from activated immune cells. SNP generates nitric oxide (NO) and reactive nitrogen species (RNS).

RNA was made from primary mouse OPCs from WT and BHMT^-/-^ using Trizol (Thermo Scientific, 15596026) and phenol:chloroform extraction. qRT-PCR was performed with gene specific primers for Pyruvate Dehydrogenase Kinase 1 (Pdk1). Primers for Pdk1 were: Forward 5’-CAGGCCCTGAGCCTTACTG-3’, Reverse 5’ AACGAGGTCTTTTCACAAGCA-3’. Relative levels of mRNA expression were determined by the 2^-ΔΔCt^ method following normalization to β-actin. Results are from three separate experiments and statistical significance was determined in GraphPad Prism with a two-way ANOVA, with p ≤ 0.05 considered significant.

### 2.8 DNMT Activity Assay

DNMT activity was quantified with an EpiQuick DNA Methyltransferase Activity/Inhibition Assay Kit (Epigentek Inc., Farmingdale, NY, P-3001-1). Primary mouse OPCs from WT and BHMT^-/-^ mice were cultured for 7 days and were then treated overnight with 1mM betaine. The following day, cells were scraped with disposable cell scrapers and pelleted via centrifugation (1500 rpm for five minutes at 4°C). Nuclear extracts were prepared as previously described (Holden & Horton, 2003). Six µg of protein from at least three separate preparations of primary cultures were measured. Statistical significance was determined in GraphPad Prism with a two-way ANOVA, p < 0.05 considered significant.

### 2.9 Respirometry

A real-time ATP rate assay was used to quantify the production of ATP from mitochondrial and glycolytic processes under conditions of stress in WT and BHMT^-/-^ primary mouse OPC cultures. Primary mouse OPCs were cultured for 7 days and were plated on PDL-coated Seahorse cell plates at a density of 12,000 cells/well. Treatments with SNP (100 uM) and betaine (1 mM) were done overnight. The next day, the media was changed to ATP basal media (DMEM [pH 7.4], 10 mM glucose, 1 mM pyruvate, 2 mM glutamine) and the cells were incubated for 1 hr at 37°C in a non-CO_2_ incubator. To run the assay, 1.5 µM oligomycin and 0.5 µM rotenone/antimycin A were injected in series to inhibit ATP synthase and mitochondrial complexes I and III, respectively. The total ATP production rate was calculated in pmol/minute/cells. Basal respiration was monitored to establish a baseline. The glycolytic and mitochondrial ATP production rates were determined by a series of biochemical equations outlined in Supplemental Figure 1. Statistical significance was determined from five separate preparations of cells in GraphPad Prism with a one-way ANOVA, p < 0.05 considered significant.

## 3. Results

### 3.1 BHMT is enriched in chromatin at genes involved in energy metabolism

ChIP-seq identified a total of 4,063 genes with statistically significant (*q*-value < 0.001) BHMT-enriched regions within a 5 kb TSS window cutoff in oligodendrocytes. Those genes included ontology categories involved in DNA binding, cellular metabolic process, and cellular components including the nucleus and membrane bound organelles (categories and numbers of genes are shown in Supplementary Fig. 1).

KEGG analysis shows that BHMT is enriched in gene regulatory regions of genes involved in hypoxia inducible factor 1 (HIF-1) signaling, glycolysis, and neurodegenerative disease (Fig. 1A). Areas enriched for BHMT include TSS, exons, and introns of genes including those involved in regulating glycolysis.

**Figure 1.**
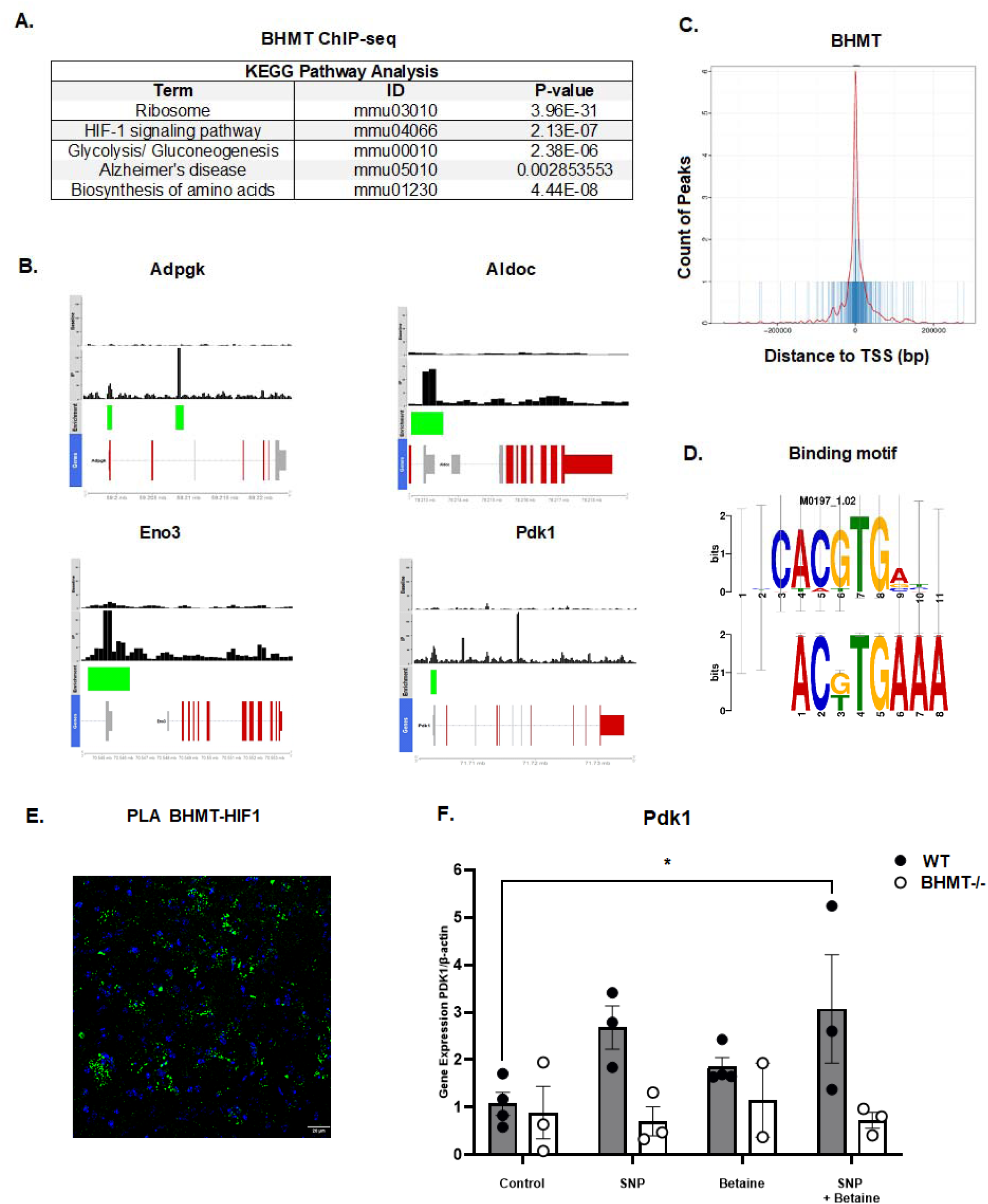
BHMT binds to chromatin at regulatory regions of genes involved in glycolysis in OPCs. **A.** ChIP-seq for BHMT in OPCs shows that BHMT is enriched in the TSS of glycolytic and maturation genes. KEGG analysis identified specific pathways that have significant enrichment with p values outlined. **B.** Enrichment for BHMT at genes involved in glycolysis including Adpgk, Aldoc, Eno3, and the Pdk1 gene are shown. Red indicates untranslated regions, blue indicates protein coding regions, yellow indicates introns. Green box indicates significant BHMT enrichment (p < 0.01). **C.** BHMT is enriched near TSS of genes. **D.** Binding motif for BHMT (bottom panel) in OPCs contains a HIF-1 binding site (ACGTG) (top panel). **E.** PLA shows that BHMT is within 40 nm of HIF1. Green fluorescence denotes the BHMT-HIF1 interaction in mouse spinal cord. **F.** Effect of betaine on *Pdk-1* expression was measured by qRT-PCR. Bar graphs illustrate betaine effects on expression of *Pdk1* in WT and BHMT^-/-^ OPCs treated with 100 µM SNP and/or 1 mM betaine relative to β-actin (n=3). Error bars presented as SEM. * p <0.05.

Further processing and visualization in R for areas of significant enrichment in glycolysis genes including the key regulator of glycolysis Pdk1 are shown in Fig. 1B. This gene is of particular interest because it is a target of HIF-1 and serves as a metabolic switch to increase glycolysis (Kim et al., 2006; Peng et al., 2018). Activation of Pdk1 would be important for OPCs as they switch to increased glycolysis during maturation to myelinating oligodendrocytes. BHMT enrichment was also found in the TSS of the ADP-dependent glucokinase (Adpgk) gene (Fig. 1B). This gene has been shown to switch metabolism to Warburg-like aerobic glycolysis in activated immune cells (Imle et al., 2029). The aldolase c (Aldoc) and enolase 3 (Eno3) genes are other enzymes of glycolysis enriched for BHMT in OPCs (Fig. 1B). Overall, BHMT enrichment was found within 2,000 bp of the TSS of genes (Fig. 1C). Motif analysis also identified a HIF-1 response element within the BHMT DNA binding sequence (Fig. 1D) suggesting that BHMT may complex with HIF-1 to regulate glycolysis. PLA analysis shows that BHMT interacts with HIF1 (within 40 nm) (Fig. 1E). Supplemental Table 1 outlines genes identified by ChIP-seq relating to metabolism and maturation that were significantly enriched and potentially regulated by BHMT.

### 3.2 BHMT regulates the expression of Pdk1 in OPCs under oxidative conditions

The transcriptional consequences of BHMT enrichment on expression of *Pdk1* was determined using qRT-PCR. Pdk1 is an enzyme responsible for mediating a metabolic switch from mitochondrial respiration to glycolysis by phosphorylation of pyruvate dehydrogenase. This gene was selected as a candidate gene to test effects of the BHMT -betaine pathway, because Pdk1 metabolically reprograms cells to increase glycolysis and is activated by HIF-1 under oxidative and inflammatory conditions (Kim et al., 2006). Cells were treated with the nitric oxide donor SNP to mimic increased RNS resulting from neuroinflammatory conditions in MS. qRT-PCR was performed in control, 100 μM SNP treated, 1 mM betaine treated, and SNP + betaine treated OPCs from WT and BHMT^-/-^ mice. Betaine treatment only increased expression of *Pdk1* mRNA in WT OPCs treated with SNP + betaine (DF=17, F=12.34, p =0.0458) indicating that the BHMT pathway is activated under inflammatory and oxidative conditions. BHMT interaction with HIF-1 on chromatin may explain this increased activation of Pdk1 with betaine treatment under oxidative conditions. Betaine effects were dependent on BHMT as there was no effect of betaine on *Pdk1* in BHMT^-/-^ OPCs (Fig 1 F).

### 3.3 BHMT regulates DNMT activity and interacts with TET1 on chromatin

To better understand the mechanism of BHMT on regulation of transcription in OPCs, we tested whether betaine could influence DNMT activity in mouse primary OPCs (Fig. 2). DNA methyltransferase activity assays were carried out on WT and BHMT^-/-^ primary mouse OPCs treated with betaine to assess BHMT-mediated changes in epigenetic function. Total DNMT activity was quantified and shows that betaine significantly increases DNMT activity (Fig. 2A) (DF=8, F=5.396, p=0.0464). This increase in DNMT activity was dependent on the presence of BHMT as demonstrated by the fact that betaine had no effect on DNMT activity in OPCs from BHMT^-/-^ mice.

**Figure 2.**
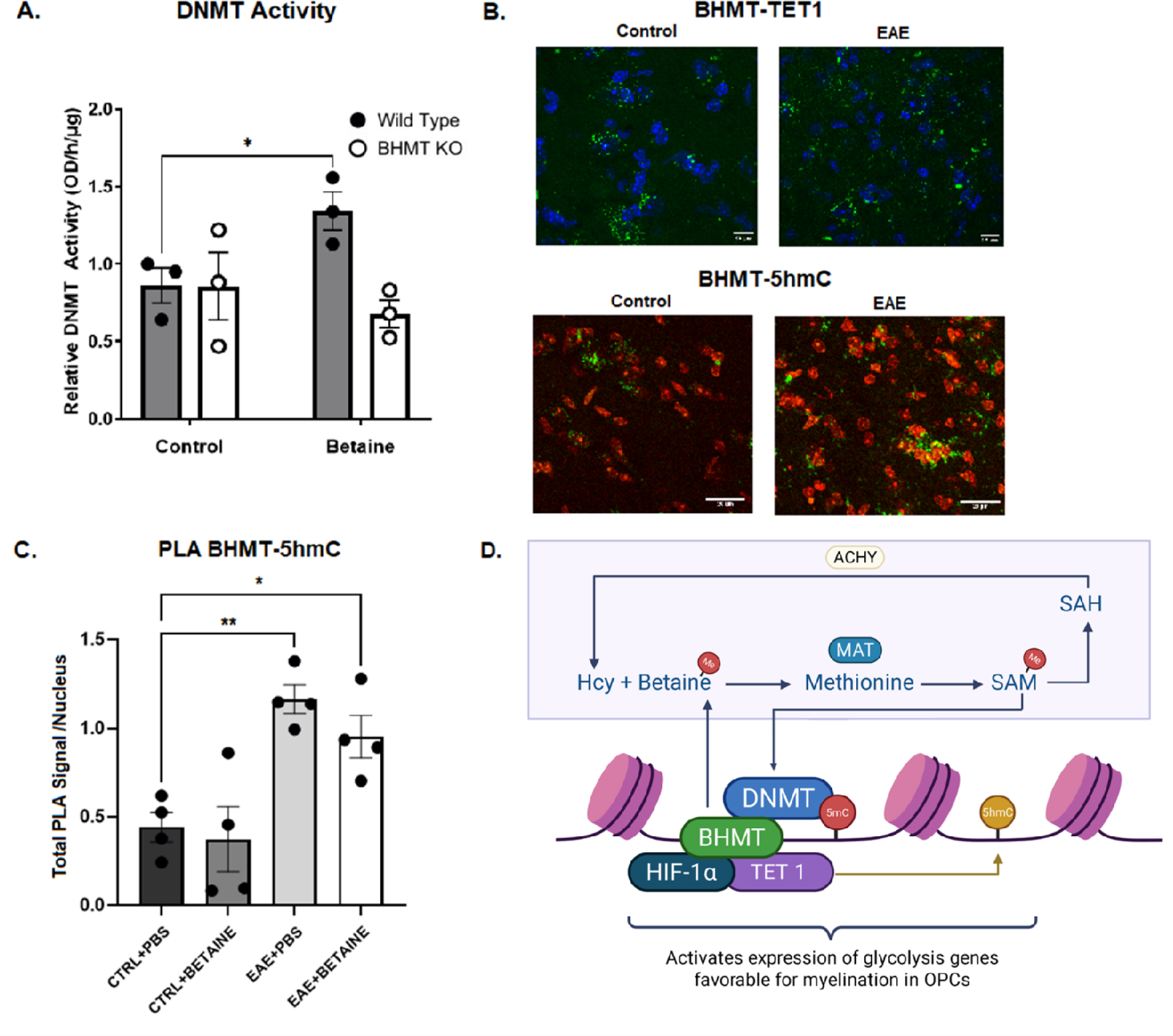
BHMT regulates methylation processes and interacts with TET1 and 5-hmC. **A.** BHMT regulates DNMT activity in OPCs. Total DNMT activity was measured in control WT and BHMT^-/-^ OPCs and in cells treated with betaine (n=3). Betaine mediated activation of DNMT depends on expression of BHMT. **B.** Representative PLA images show that BHMT interacts with TET1 and 5-hmC in control and EAE mouse lumbar spinal cord. Green fluorescence denotes a BHMT and TET1 or 5-hmC interaction in naïve mice and in EAE mice. Nuclei are visualized with DAPI (blue or red). **C.** Quantitation of the PLA signal shows increased BHMT-5hmC interaction (n=4) in EAE lumbar spinal cord compared to control. **D.** Schematic depicts BHMT interactions with DNMTs and TET1 on chromatin that regulate methylation and hydroxymethylation and transcription of genes necessary for OPC maturation and remyelination. Betaine is the methyl donor for BHMT to remethylate homocysteine to methionine. The MAT enzyme then generates SAM from methionine. SAM is the methyl donor for DNMT catalyzed DNA methylation that can subsequently be partially demethylated to 5-hmC by TET1. Error bars represent SEM. * p < 0.05, ** p < 0.01.

We have previously shown that BHMT interacts with DNMTs (Sternbach et al., 2021). A proximal ligation assay shows that BHMT also interacts with TET1 and 5-hmC in mouse lumbar spinal cord (Fig. 2 B). These data show that BHMT is within 40 nm of TET1 and 5-hmC on chromatin. Quantitation of PLA signals shows that the BHMT-5hmC interaction is significantly increased in EAE mice with or without betaine (Fig. 2C) (DF=12, F=9.709, *p = 0.013, **p = 0.0014). Taken together, BHMT interactions with DNMT, TET1, and 5-hmC suggests that the BHMT -betaine pathway can regulate DNMT activity and 5-mC at specific sites that are subsequently converted to 5-hmC to activate transcription to support OPC metabolism and enhance remyelination (Fig 2 D). Further studies will be required to identify specific loci with enhanced 5-hmC in oligodendrocytes with betaine mediated activation of the BHMT pathway.

### 3.4 The BHMT -betaine pathway regulates glycolysis in OPCs

Our ChIP-seq data indicated that the BHMT enzyme was bound to promoter regions of genes involved in OPC energy metabolism. To assess the functional consequences of this relationship we investigated the effect of the absence of the BHMT enzyme on epigenetic control of metabolism under conditions of oxidative stress. To address the downstream implications of the loss of the BHMT -betaine pathway on OPC energy metabolism, we performed Seahorse Real-Time ATP Production Rate Assays to quantitate the relative contribution of ATP production from glycolysis and oxidative phosphorylation in WT and BHMT^-/-^ OPCs. These data show that WT OPCs synthesize ATP by a combination of oxidative phosphorylation (mito) and glycolysis (approximately 50% glycolysis to 50% oxidative phosphorylation) (Fig. 3A, B, and C) while OPCs isolated from BHMT^-/-^ mice show impaired glycolysis and high levels of oxidative phosphorylation (20% glycolysis to almost 80% oxidative phosphorylation) (Fig. 3D, E, and F). We then examined the effects of oxidative conditions on OPC metabolism by treating cells with SNP. With SNP treatment, ATP production was severely compromised in WT and in BHMT^-/-^ cells. SNP treatment significantly inhibited glycolysis in WT OPCs. The effects of betaine on repairing OPC energetics under oxidative conditions was then tested. In WT OPCs treated with SNP, betaine restored ATP levels back to control levels by increasing glycolysis (Fig. 3A, B) (DF=16, F=2.610, p = 0.0209 for SNP treated cells, p = 0.0390 for SNP + betaine treated cells). However, betaine had no effect on oxidative phosphorylation (Fig. 3C). Treatment with betaine had no significant effect on either glycolytic or oxidative ATP production in BHMT^-/-^ OPCs (Fig. 3D, E, and F).

**Figure 3.**
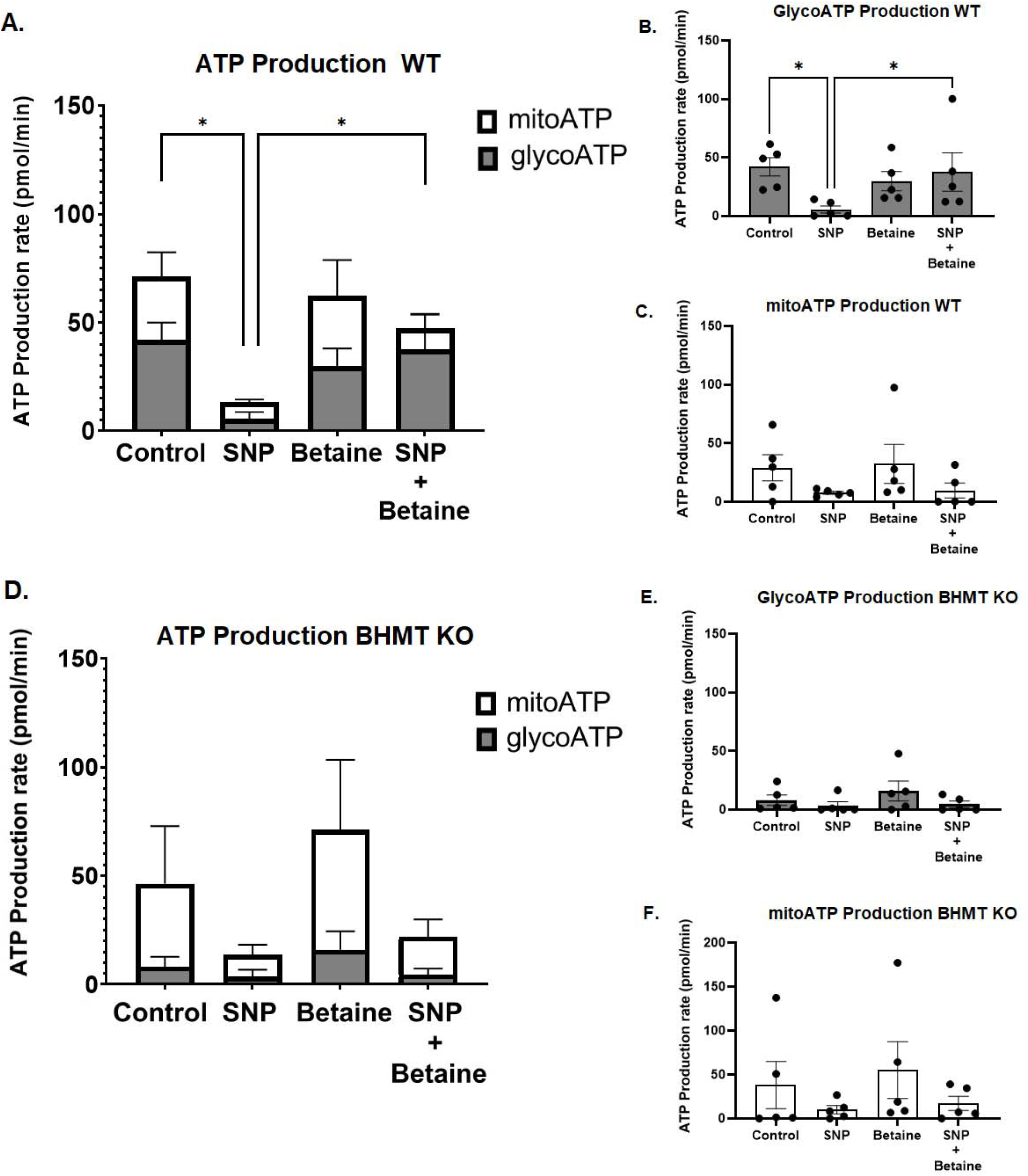
Seahorse respirometry shows that BHMT regulates glycolysis in OPCs. **A.** The contribution of glycolysis and oxidative phosphorylation (mito) to the total ATP production rate is shown for WT OPCs treated with 100 µM SNP and/or 1 mM betaine (n=5). **In B and C,** the contribution of ATP production by glycolysis and oxidative phosphorylation for WT OPCs are shown separately. **D.** Total ATP production rate for glycolysis and oxidative phosphorylation in BHMT knockout OPCs treated with 100 µM SNP and/or 1 mM betaine (n=5). **E and F** show ATP production rates for glycolysis and oxidative phosphorylation separately for BHMT knockout OPCs (n=5). Error bars denote SEM. * p < 0.05.

### 3.5 Betaine treated EAE mice exhibit improved clinical scores and increased myelin and neuron survival in spinal cord

EAE mice receiving betaine showed significantly improved clinical scores (Fig. 4 A) in acute and especially in chronic disease stages (day 25-40 post-immunization) (DF=52, T=4.616, p = 0.0228). Mean disability scores improved from 3.4 in EAE mice in chronic stages to 2.5 in mice treated with betaine from day 15 to day 40. IHC for NeuN+ neurons shows that betaine treatment is neuroprotective. EAE mice treated with betaine showed a significant improvement in neuron survival compared to EAE mice given PBS (Fig. 4 B, C). EAE mice had about 50% fewer neurons in the ventral horn compared to control mice, while EAE mice treated with betaine showed no significant change in the number of NeuN+ neurons from controls (DF=3, 12; F=10.77; *** p = 0.0005, * p = 0.030). IHC of lumbar spinal cord sections with an antibody to MBP also revealed increased myelin protein in EAE mice treated with betaine (Fig. 4 D, E) (DF=8, F=13.09, p = 0.0068). MBP levels were enhanced approximately three-fold in lumbar spinal cord of mice receiving betaine. CARS showed that myelin lipids are also significantly increased by three-fold in lumbar spinal cord of EAE mice treated with betaine (Fig. 4 F, G) (DF=5, t=6.312, p = 0.0015).

**Figure 4.**
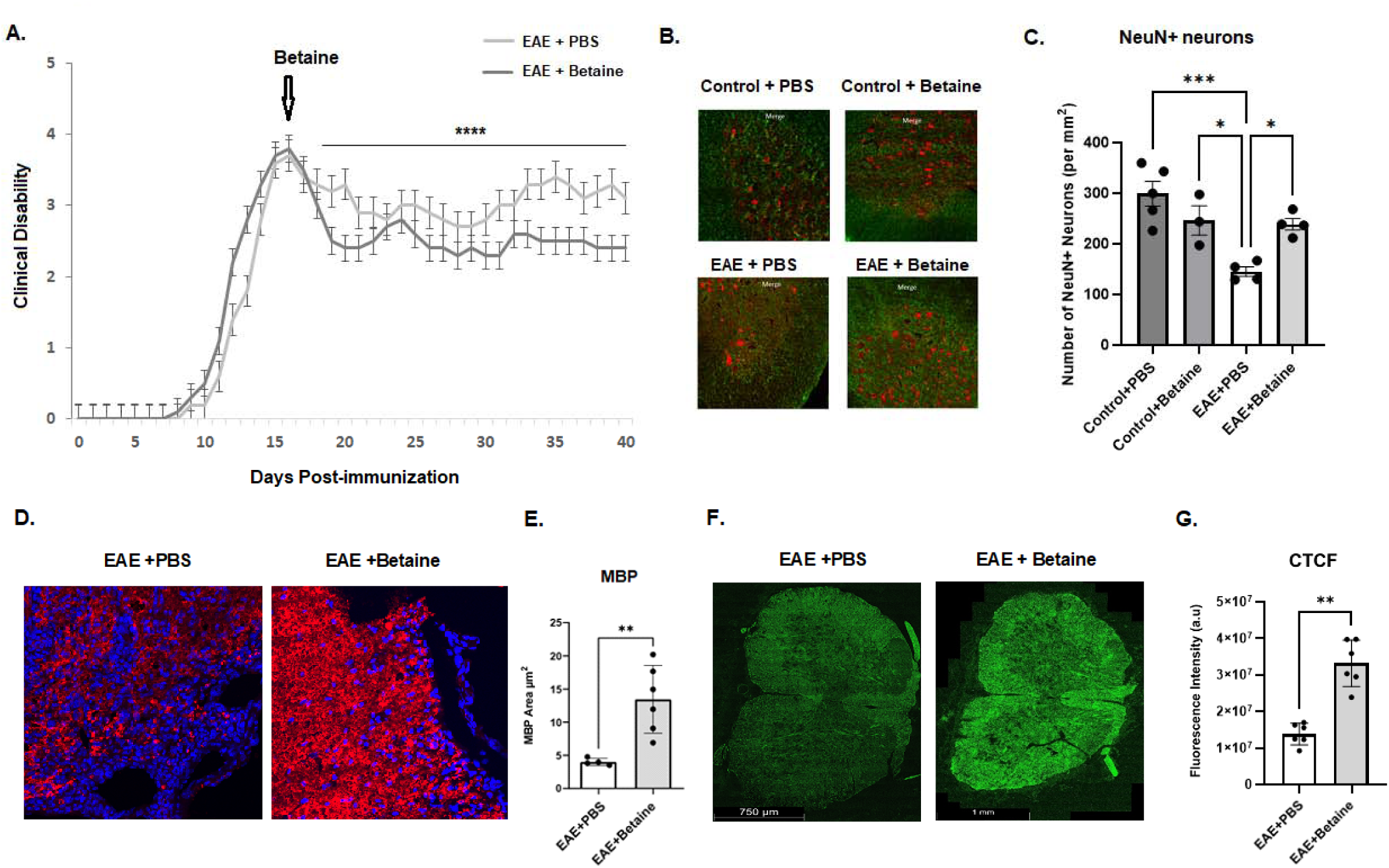
Betaine provides neuroprotection and enhances myelin in EAE. Mice were given ip injections of betaine or vehicle (PBS) starting at day 15 post-immunization. **A.** Betaine significantly improved clinical scores in EAE mice (n=10). **B.** Representative confocal images of NeuN+ (red fluorescence) in ventral motor neurons and MBP (green fluorescence) in lumbar spinal cord of control and EAE mice given either betaine or PBS injections and sacrificed at day 40 post-immunization. **C.** Quantitation shows no significant difference in NeuN+ neurons/mm^2^ between control mice treated with betaine (n=3) or PBS (n=5), however, betaine treatment increased numbers of NeuN+ ventral motor neurons in lumbar spinal cord of EAE mice compared to control EAE mice given PBS (n=4). **D.** Representative IHC images for MBP (red) and DAPI (blue) on lumbar spinal cord sections from EAE and EAE + betaine mice. **E.** Quantitation for MBP shows that EAE mice treated with betaine (n=6) exhibited reduced lesion area compared to EAE mice treated with vehicle (n=4). **F.** Representative CARS image shows reduced lipid fluorescence in EAE spinal cord compared to EAE + betaine treated mice. **G.** Quantitation of corrected total cellular lipid fluorescence (CTCF) from lumbar spinal cords of EAE mice treated with vehicle or betaine (n=6). Fluorescence was measured in nine lumbar spinal cord sections from six mice in each group. Error bars represent SEM. * p <0.05, ** p < 0.01, *** p < 0.005, **** p < 0.001.

## 4. Discussion

While other studies have reported protective effects of betaine in the periphery on inflammation and aging, (Geng et al., 2025) these effects don’t appear to involve BHMT. Centrally, in the nervous system, we show that betaine’s epigenetic effects are mediated through BHMT mediated transmethylation reactions, supplying SAM locally on chromatin to activate transcriptional programs in OPCs to enhance myelination. In support, earlier studies indicated that betaine treatment significantly increased total DNMT activity in human MO3.13 oligodendrocytes, while this effect was not seen when betaine treatment was accompanied by siRNA knockdown of BHMT (Sternbach et al., 2021). Mechanistically, in the present study, we show that the effects of betaine depend on the presence of BHMT as betaine shows no effect on DNMT activity, expression of key regulators of glycolysis, or ATP production in OPCs isolated from BHMT^-/-^ mice.

We have shown that the BHMT methylation pathway regulates OPC metabolism to increase glycolysis by activating Pdk1 and glycolysis in oxidative and inflammatory states, providing neuroprotection and enhanced myelin in EAE. Our data demonstrate the importance of one carbon metabolism, specifically the BHMT-betaine pathway, to oligodendrocyte metabolism and myelin stability. Earlier studies from our laboratory found that concentrations of one carbon cycle metabolites, including betaine and the methyl donor SAM, were significantly decreased in MS postmortem brain tissue (Singhal et al., 2015). One carbon metabolites were also found to be altered peripherally in sera from individuals with early-stage MS (Singhal et al., 2018). Given that one carbon metabolism was dysregulated in MS, we reasoned that there were downstream epigenetic consequences. We confirmed this in neurons and found that betaine epigenetically regulates mitochondrial genes involved in oxidative phosphorylation by increasing H3K4me3 (Singhal et al., 2015; 2020). We also found that murine OPCs and oligodendrocytes express BHMT in the cytoplasm and nucleus, as well as human OPCs in sclerotic lesions (Sternbach et al., 2021). This is critical as lesions are areas devoid of myelin, and it has been reported that progenitor cells will migrate to sites of lesions and fail to differentiate (Kuhlmann et al., 2008). The presence of OPCs expressing BHMT in lesions indicates the potential for epigenetic intervention to tip the scales in favor of OPC maturation and remyelination with betaine administration.

Previous studies have focused on the dynamics of the oligodendrocyte lineage and how epigenetics can influence maturation (Egawa et al., 2019; Emery & Lu, 2015; Liu et al., 2015; Moyon & Casaccia, 2017). However, few have identified underlying epigenetic mechanisms that fail under conditions of oxidative stress, and fewer have revealed a means for repair (Spaas et al., 2021). Our lab has previously shown that betaine restores levels of the methyl donor SAM and improves epigenetic control in the cuprizone mouse model of MS (Singhal et al., 2015; 2020). In the present study we examine the role of betaine and the BHMT-betaine pathway in the inflammatory EAE model of MS. Our data show that this pathway increases methyltransferase activity, shifts OPC metabolism, and provides protection in EAE.

ChIP-seq analysis of BHMT in primary OPC cultures found significant BHMT enrichment at over 60 genes which pertained to differentiation and metabolic processes. Specifically, BHMT was enriched at Sox10 and Tcf7l2 loci, two critical regulators of differentiation that are also necessary for efficient remyelination processes (Hornig et al., 2013; Lopez-Anido et al., 2015; Wegener et al., 2015; Guo et al., 2023). This is consistent with our previous study showing that betaine upregulates Sox10 under oxidative conditions in cultured OPCs (Sternbach et al., 2021). Moreover, BHMT was also enriched at genes involved in glycolysis including Adpgk, Aldoc, Eno3, and Pdk1. This enrichment further suggests a role for the BHMT-betaine methylation pathway in local epigenetic modulation of OPC metabolism. Interestingly, BHMT was enriched in KEGG pathways of glycolysis and HIF-1 signaling.

Pdk-1 is a critical regulator of glycolysis and is activated by HIF-1. We found the betaine increases expression of Pdk-1 under oxidative conditions, indicating that the BHMT -betaine pathway may act with HIF-1 to activate glycolysis under inflammatory or oxidative states.

We show that BHMT is enriched at genes that pertain to maturation and metabolism in OPCs. Activating DNMTs and DNA methylation at these genomic loci would be expected to silence transcription of associated genes, however we observe that BHMT is enriched at the TSS of Pdk1 and that betaine upregulates transcription of the mRNA for Pdk1. This can be explained by another key player in this modulation which is DNA hydroxymethylation (5-hmC), an epigenetic modification, initially mediated by DNMT, then followed by oxidation of 5-mC to 5-hmC by TET enzymes. DNA hydroxymethylation is associated with transcriptional activation and TET1 has been shown to be important for remyelination by adult OPCs (Moyon et al., 2021). Our data showing that BHMT interacts with TET1 and is in close proximity (within 40 nm) of 5-hmC suggests that the BHMT-betaine pathway can both activate and repress gene transcription through DNMT activities.

Our data are consistent with other studies that have shown a link between oligodendrocyte energy metabolism and myelin. OPC maturation is dependent upon cellular energetics (Jeffries et al., 2015; Cabello-Rivera et al., 2019) and maturing oligodendrocytes preferentially use glycolysis for ATP production (Rao et al., 2017). However, mature oligodendrocytes will switch to oxidative phosphorylation when under stress, resulting in impaired myelin generation and maintenance as a means of survival (Harris & Atwell, 2012; Rone et al., 2016). We report here that oxidative stress significantly decreased total ATP production in OPCs, and that this can be remedied with betaine through activation of the glycolytic pathway. The rescue effect of betaine on OPC energy metabolism is supported by our ChIP-seq data that identified a HIF-1 binding site in the BHMT DNA binding motif and indicated significant enrichment of BHMT in genes involved in glycolysis, including Pdk1, a key player in the molecular switch from oxidative phosphorylation to glycolysis (Peng et al., 2018). This change in OPC metabolism elicited by betaine has functional consequences as shown in our EAE studies. In EAE mice supplemented with betaine at the peak of clinical disability, motor ability was improved and lesion area was decreased. Betaine improved clinical scores most significantly during the chronic phase of disease from day 25 to day 40 post-injection indicating neuroprotection. Consistent with a role in overcoming the maturation block of OPCs, betaine increased myelin protein and myelin lipids in the inflammatory EAE mouse model, suggesting that betaine enhances OPC maturation.

Overall, these data support and expand upon previous findings showing that activating the BHMT-betaine pathway enables epigenetic modulation in an MS-like environment (Singhal et al., 2020). Taken together, our respirometry data showing that the BHMT-betaine pathway plays a role in switching OPC metabolism to increase glycolysis and our CARS data showing that betaine treatment increases myelin lipids suggests that the metabolic switch is increasing acetyl-CoA required to support myelination under oxidative or inflammatory conditions. These studies have expanded our knowledge on BHMT’s role in the CNS and how enhancing methionine metabolism with betaine can attenuate disease.

## 5. Conclusions

Here, we have outlined the role of the BHMT-betaine pathway in oligodendrocytes and how these epigenetic modifiers contribute to protection in EAE. This study is the first to report that BHMT mediated methylation influences epigenetic control of OPC energy metabolism. Limitations of these findings are that the genes regulated by methylation or hydroxymethylation remain to be identified in future studies. It also isn’t clear whether the BHMT-betaine pathway is protecting myelin or supporting remyelination in EAE. Further studies need to be done to distinguish between these possibilities.

However, results from this study are clinically relevant as they suggest that epigenetic therapies will be efficacious in managing demyelinating and neurodegenerative disorders, including MS, Alzheimer’s disease, and Parkinson’s disease.

## Acknowledgements

This work was funded in part by a pilot grant from the BHRI at Kent State University (JM, JLW) and from a grant from the Department of Biological Sciences at Kent State University (JM).

## Author Contributions

SS and JM conceived of the study and wrote the manuscript. SS, MWP, NAR, MSM, KK collected and analyzed data, AE performed data analysis for ChIP-seq, RC, EJF, and JLW assisted with data collection and analyzed data, SZ donated BHMT knockout mice and contributed to data collection.

## Conflicts of Interest

The authors have no conflict of interest related to the data presented in this study.

## Abbreviations

Adpgk: ADP-Dependent Glucokinase
Aldoc: Aldolase C
ATP: Adenosine Triphosphate
BHMT: Betaine-Homocysteine Methyltransferase
BHMT^-/-^: Betaine Homocysteine Methyltransferase Knockout Mice
CARS: Coherent Anti-Stokes Raman Scattering
ChIP-seq: Chromatin Immunoprecipitation Sequencing
CNS: Central Nervous System
CTCF: Corrected Total Cell Fluorescence
DAPI: 4,6-diamidino-2-phenylindole
DNA: Deoxyribonucleic Acid
DNMT: DNA Methyltransferase
EAE: Experimental Autoimmune Encephalomyelitis
Eno3: Enolase 3
H3K4me3: Histone 3 Lysine 4 Trimethylation
HIF-1: Hypoxia inducible factor 1
Hk1: Hexokinase 1
IACUC: Institutional Animal Care and Use Committee
IHC: Immunohistochemistry
KEGG: Kyoto Encyclopedia of Genes and Genomes
MBP: Myelin Basic Protein
MS: Multiple Sclerosis
NA: Numerical Aperture
OPCs: Oligodendrocyte Precursor Cells
OLs: Myelin-making Oligodendrocytes
PBS: Phosphate-Buffered Saline
Pdk1: Pyruvate Dehydrogenase Kinase 1
PFA: Paraformaldehyde
PLA: Proximal Ligation Assay
qRT-PCR: Quantitative Real-Time Polymerase Chain Reaction
RNA: Ribonucleic acid
RNS: Reactive Nitrogen Species
ROI: Region of Interest
RRID: Research Resource Identifier
SAM: S-adenosyl Methionine
SHG: Second Harmonic Generation
SNP: Sodium Nitroprusside
TET1: Ten-Eleven Translocation 1
TSS: Transcription Start Site
WT: Wild-Type
5-hmC: 5-Hydroxymethylcytosine
5-mC: 5-Methylcytosine

## Notes

### Competing Interest Statement

The authors have declared no competing interest.

