## Supplemental Figures for "The BHMT-TET1 axis regulates glycolytic metabolism in oligodendrocytes and increases myelin in the EAE mouse model of multiple sclerosis"

**Supplemental Figure 1**

**Equations for calculation of total ATP production rate**

Glycolytic ATP production rate (GlycoATP production rate):

Eq. 1. Glucose + 2 ADP + 2 Pi 🡪 2 Lactate + 2 ATP + 2 H_2_0 + 2 H^+^

ATP production in Eq. 1. is the same as the glycolytic protein efflux rate (GlycoPER), thus:

Eq. 2. GlycoATP production rate (pmol ATP/min) = GlycoPER (pmol H^+^/min)

Mitochondrial ATP production rate (mitoATP production rate):

The oxygen consumption rate (OCR) coupled to ATP production during oxidative phosphorylation is calculated as the OCR that is inhibited by oligomycin, thus:

Eq. 3. OCR (pmol O_2_/min) - OCR_Oligo_ (pmol O_2_/min) = OCR_ATP_ (pmol O_2_/min)

Transformation of OCR_ATP_ to match the rate of mitochondrial ATP production:

Eq. 4. OCR_ATP_ (pmol O_2_/min) * 2 (pmol O/pmol O_2_) * P/O (pmol ATP/mol O) = mitoATP production rate (pmol ATP/min)

where P/O is the number of ADP molecules phosphorylated to ATP per oxygen (O) atom, 2.75

Total cellular ATP production rate is the sum of glycolytic and mitochondrial ATP production rates:

Eq. 5. glycoATP production rate (pmol ATP/min) + mitoATP production rate (pmol ATP/min) = ATP production rate (pmol ATP/min)

**Supplementary Figure 2**

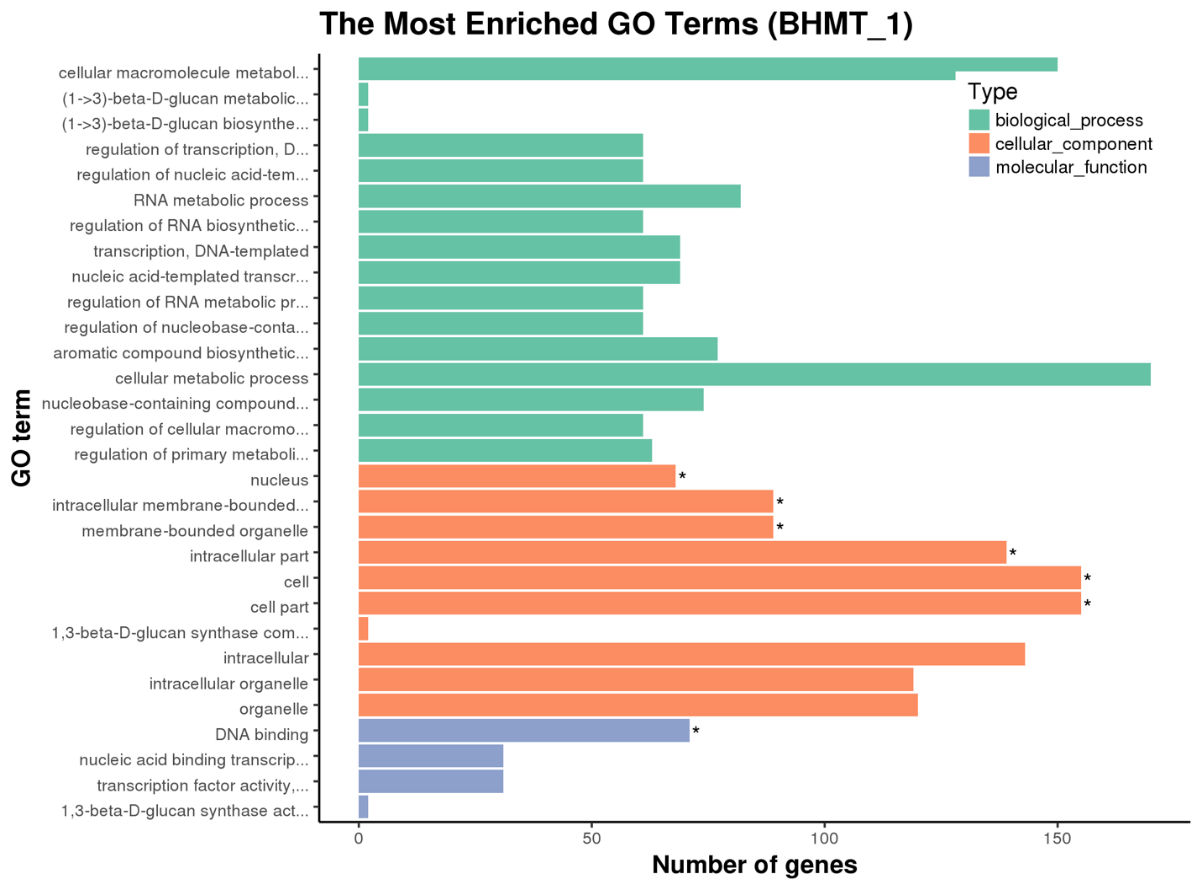

**Supplementary Table 1. BHMT enrichment in metabolism and maturation genes.**

| **Chromosome** | **Gene** | **Start_Position** | **End_Position** | **Strand** | **TSS** |
| --- | --- | --- | --- | --- | --- |
| 1 | Sox13 | 133310041 | 133352115 | -1 | 133352115 |
| 2 | Abca2 | 25318715 | 25338552 | 1 | 25318715 |
| 2 | Pdk1 | 71703568 | 71734202 | 1 | 71703568 |
| 2 | Ptpra | 130292198 | 130398044 | 1 | 130292198 |
| 2 | Sox12 | 152235531 | 152239983 | -1 | 152239983 |
| 2 | Pfkfb3 | 11476244 | 11558888 | -1 | 11558888 |
| 3 | Med12l | 58913246 | 59226103 | 1 | 58913246 |
| 4 | Trp73 | 154140706 | 154224665 | -1 | 154224665 |
| 4 | Chchd2-ps | 148152488 | 148152949 | -1 | 148152949 |
| 4 | Galt | 41755228 | 41758695 | 1 | 41755228 |
| 4 | Pink1 | 138040720 | 138053618 | -1 | 138053618 |
| 4 | Slc25a33 | 149828493 | 149858734 | -1 | 149858734 |
| 5 | Cdk5 | 24623239 | 24628528 | -1 | 24628528 |
| 5 | Fgfr3 | 33879018 | 33894412 | 1 | 33879018 |
| 5 | Nkx6-1 | 101806005 | 101812862 | -1 | 101812862 |
| 5 | Rheb | 25007821 | 25047622 | -1 | 25047622 |
| 5 | Wdr1 | 38684156 | 38720564 | -1 | 38720564 |
| 5 | Foxk1 | 142387252 | 142447766 | 1 | 142387252 |
| 5 | Iscu | 113910809 | 113916349 | 1 | 113910809 |
| 5 | Slc4a4 | 89034345 | 89387512 | 1 | 89034345 |
| 6 | Pparg | 115337912 | 115467360 | 1 | 115337912 |
| 8 | Nrg1 | 32304579 | 33374825 | -1 | 33374825 |
| 8 | Sox1 | 12445295 | 12450126 | 1 | 12445295 |
| 8 | Cox4i1 | 121394961 | 121400946 | 1 | 121394961 |
| 9 | Dag1 | 108081833 | 108141157 | -1 | 108141157 |
| 9 | Adpgk | 59198841 | 59231335 | 1 | 59198841 |
| 9 | Rhoa | 108183328 | 108215133 | 1 | 108183328 |
| 10 | Ascl1 | 87326681 | 87329522 | -1 | 87329522 |
| 10 | Hdac2 | 36850540 | 36877885 | 1 | 36850540 |
| 10 | Atp5d | 79974466 | 79981652 | 1 | 79974466 |
| 10 | Zbtb7a | 80971054 | 80988829 | 1 | 80971054 |
| 11 | Nf1 | 79230519 | 79472438 | 1 | 79230519 |
| 11 | Aldoc | 78213794 | 78218607 | 1 | 78213794 |
| 11 | Atp5h | 115306515 | 115310788 | -1 | 115310788 |
| 11 | Eno3 | 70548028 | 70553339 | 1 | 70548028 |
| 11 | Foxk2 | 121150816 | 121200722 | 1 | 121150816 |
| 11 | Git1 | 77384388 | 77398612 | 1 | 77384388 |
| 11 | Ncor1 | 62207252 | 62349367 | -1 | 62349367 |
| 12 | Dicer1 | 104654001 | 104718211 | -1 | 104718211 |
| 12 | Nkx2-1 | 56578743 | 56583693 | -1 | 56583693 |
| 12 | Dld | 31381276 | 31401452 | -1 | 31401452 |
| 12 | Entpd5 | 84420631 | 84455803 | -1 | 84455803 |
| 13 | Gli3 | 15637820 | 15904611 | 1 | 15637820 |
| 13 | Id4 | 48414704 | 48419502 | 1 | 48414704 |
| 13 | Prxl2c | 64423094 | 64460524 | -1 | 64460524 |
| 15 | Sox10 | 79039108 | 79049440 | -1 | 79049440 |
| 15 | Ep300 | 81469552 | 81536278 | 1 | 81469552 |
| 15 | Ppara | 85619184 | 85687020 | 1 | 85619184 |
| 15 | Prkaa1 | 5173343 | 5211380 | 1 | 5173343 |
| 16 | Hes1 | 29883202 | 29886614 | 1 | 29883202 |
| 16 | Olig1 | 91066660 | 91068821 | 1 | 91066660 |
| 16 | Olig2 | 91022345 | 91025565 | 1 | 91022345 |
| 16 | App | 84746573 | 84970654 | -1 | 84970654 |
| 16 | Atp5j | 84624754 | 84632513 | -1 | 84632513 |
| 16 | Zbtb20 | 42696244 | 43462965 | 1 | 42696244 |
| 17 | Mir219a-1 | 34243957 | 34244066 | -1 | 34244066 |
| 17 | Qk | 10421530 | 10538783 | -1 | 10538783 |
| 18 | Atp5a1 | 77861429 | 77870569 | 1 | 77861429 |
| 19 | Grk2 | 4336029 | 4356250 | -1 | 4356250 |
| 19 | Tcf7l2 | 55730252 | 55922086 | 1 | 55730252 |
| 19 | Fxn | 24238817 | 24257969 | -1 | 24257969 |
| 19 | Sdhaf2 | 10477898 | 10503506 | -1 | 10503506 |
